# C-di-GMP is Required for Swarming in *E. coli*: a Role for DgcO and Colanic Acid

**DOI:** 10.1101/2025.01.04.631324

**Authors:** YuneSahng Hwang, Marta Perez, Rebecca Holzel, Rasika M. Harshey

## Abstract

Many bacteria use flagella to swim individually through bulk liquid, or swarm collectively over a semi-solid surface. In *E. coli*, c-di-GMP inhibits swimming via the effector protein YcgR. We show in this study that contrary to its effect on swimming, a certain threshold level of c-di-GMP is required for swarming. Gene expression profiles first indicated that several c-di-GMP synthases – *dgcJ, dgcM, dgcO* - were upregulated during swarming. Of these, we found DgcO to play a critical role. DgcO is reported to affect PGA (poly-β-1,6-N-acetylglucosamine) synthesis in *E. coli*. We show that DgcO also promotes production of colanic acid - one of the three major exopolysaccharides in *E. coli*, and that colanic acid has hitherto-unknown surfactant properties that are expected to aid swarming.

**Importance:** It is well-established that in bacteria, c-di-GMP inhibits flagella-driven motility at various points in the pathway. Concomitantly, elevated c-di-GMP levels induce the expression and synthesis of a variety of exopolysaccharides that enmesh the bacteria in a biofilm, thereby also interfering with flagella function. This study reports the surprising finding that in *E. coli*, the exopolysaccharide colanic acid is required to enable surface navigation and that the diguanylate cyclase DgcO is employed for this purpose. For surface navigation, there appears to be a sweet spot where c-di-GMP levels are just right to produce polysaccharides that can serve as surfactants and wetting agents rather than promote formation of biofilms.

## Introduction

c-di-GMP signaling offers bacteria lifestyle choices, the most prevalent being a choice between moving freely or settling in a biofilm [1–3]. *E. coli* has multiple diguanylate cyclases (DGCs) and phosphodiesterases (PDEs) that synthesize and degrade c-di-GMP, respectively [4]. In free-swimming or planktonic *E. coli*, DgcN, DgcO, DgcQ and DgcE are reported to be some of the most active DGCs, and PdeH the most active PDE [5–7]. Elevated c-di-GMP levels resulting from the inactivation of PdeH inhibit swimming but have only a moderate effect on swarming [8]. The inhibitory effect of c-di-GMP on swimming is well-studied and is orchestrated via interaction of the effector protein YcgR with both the flagellar motor and rotor [5, 8–13]. High levels of c-di-GMP promote biofilm formation [1, 3, 14]. *E. coli* biofilms consist of various extracellular polymers and polysaccharides [15], including colanic acid (CA), cellulose, curli fimbriae, and poly-β-1,6-N-acetyl-d-glucosamine (PGA) [2].

Swarming presents unique challenges not encountered while swimming [16, 17]. These include limited water availability necessary for flagella to work, and formidable physical forces such as surface tension, friction, capillary and viscous forces. Bacteria overcome these challenges by secreting surfactants to reduce surface tension and osmolytes/wetting agents to draw water to the surface, altering cell shape to improve side-by-side alignment to lower viscous drag, and increasing flagella numbers or recruiting special stator-associated proteins to enhance motor power [17–20]. In the laboratory, swarming is observed on media solidified with agar, whose consistency can range from semi-solid to solid depending on the agar concentration (0.5%-0.7% or 1-2% w/v), presenting differing levels of navigational challenge. Bacteria that only swarm on the lower range of agar concentration have been classified as ‘temperate’, and those that swarm on the higher range as ‘robust’ swarmers [17]. *E. coli* and *Salmonella* swarmers belong to the former category, while *Proteus mirabilis* and *Vibrio parahaemolyticus* fall under the robust category; the latter set of bacteria increase flagella production to enhance motor power. Some temperate swarmers like *Serratia marcescens* and *Bacillus subtilis* secrete copious amounts of surfactants and/or increase flagella numbers [19].

*E. coli* and *Salmonella* neither increase flagella numbers nor are reported to secrete specific surfactants [21–24]. These bacteria require nutrient-rich media for swarming, with *E. coli* being particularly fastidious, needing in addition 0.5% glucose and a specific type of agar (Eiken agar) [24, 25]. Several studies suggested that lipopolysaccharide (LPS) might serve to attract water to the surface in these bacteria and implicated an additional role for flagella rotation in enabling a wet surface [26–29]. One of these studies identified non-swarming mutants in the LPS biosynthetic pathway of *Salmonella* and suggested that the hydrophilic nature of LPS may serve as a wetting agent [25], while another detected high osmotic pressure at the leading edge of an *E. coli* swarm using fluorescent liposomes, and suggested that a secreted high molecular weight substance (perhaps LPS) extracts water from the underlying agar to enable motility [26]. Microarray data showed that LPS synthesis was upregulated specifically in *Salmonella* swarms [30]. Non-swarming mutants of *Salmonella* also mapped to the CA biosynthesis pathway [26]. CA is a polyanionic heteropolysaccharide composed of repeating hexo-saccharides and could also serve as a potential secreted wetting agent. We note that in the surfactant literature, the terms ‘surfactant’ and ‘wetting agent’ have been used interchangeably; here we make a distinction and use wetting agent to mean ‘ability to attract water’.

The present study was initiated to understand the significance of the differential expression of several DGCs and PDEs observed in RNAseq data collected during *E. coli* swarming. Of the upregulated DGCs, DgcJ is reported to associate with and activate NfrB glycosyl transferase to produce an exopolysaccharide that is a receptor for phage N4 [31, 32], DgcM is reported to regulate the production of curli fibers through the *csgABC* operon [33–35], and DgcO is recognized for its ability to bind oxygen via its heme domain [36] and to enhance PGA synthesis [37]. Deletion analysis of the upregulated genes indicated that only absence of DgcM and DgcO had a negative effect on swarming, the absence of DgcO being critical.

We therefore investigated the role of DgcO and found that its swarming defect is caused primarily by a lack of CA production. We demonstrate that CA has surfactant properties that, along with its expected wetting properties, assist swarming.

## Results

### DgcO plays a critical role in swarming

Mutations in *dgcJ* were recovered during a screen for motile antibiotic-resistant mutants emerging from the edge of an *E. coli* swarm, suggesting that DgcJ is active in the swarm [38]. That c-di-GMP enzymes might be differentially regulated during swimming vs swarming was inferred by the differential effect of the absence of PdeH on the two types of motilities [8] (Figure S1A). Estimation of c-di-GMP levels in WT under the two conditions using a riboswitch-based c-di-GMP sensor [39], showed these to be similar (Table 1). However, there was a much larger increase of c-di-GMP in the Δ*pdeH* mutant in swim compared to swarm cells (Fig. S1B and Table 1). These observations motivated us to examine RNAseq data (generated as part of a different project; see Methods) for expression of DGCs and PDEs during a 2-20 h the time course of swarming. Compared to planktonic or swim cells, we observed that the expression of *dgcJ, dgcM, dgcO* was elevated ∼3-8 fold during this time course (Fig. 1A). Among the PDEs, *pdeH* was drastically downregulated during swarming, with a reduction of over 100-fold in the measured raw values (Fig. S1C), while *pdeO* and *pdeR* were significantly upregulated (Fig. S1C). The contribution of the upregulated DGCs and PDEs to both swimming swarming was investigated next by deleting them individually (Fig. 1B,C and Fig. S1D). Deletion of neither *pdeO* nor *pdeR* had any effect on either motility (Fig. S1D). Deletion of *dgcJ* also had no observable effect, deletion of *dgcM* decreased swarming (an effect more evident if the plates were allowed to dry longer prior to inoculation), while deletion of *dgcO* had a consistently worse outcome for swarming (Fig. 1C). When Δ*dgcO* cells were observed under the microscope at 4h, a time point when active motion is observable in WT, they were seen to be moving as well as WT (Movies S1 & S2), showing that the defect was not in motility, but in their ability to advance across the surface. We conclude that both DgcM and DgcO play a positive role specifically during swarming, with DgcO being the dominant player. Going forward, we focused on the contribution of DgcO.

**Figure 1.**
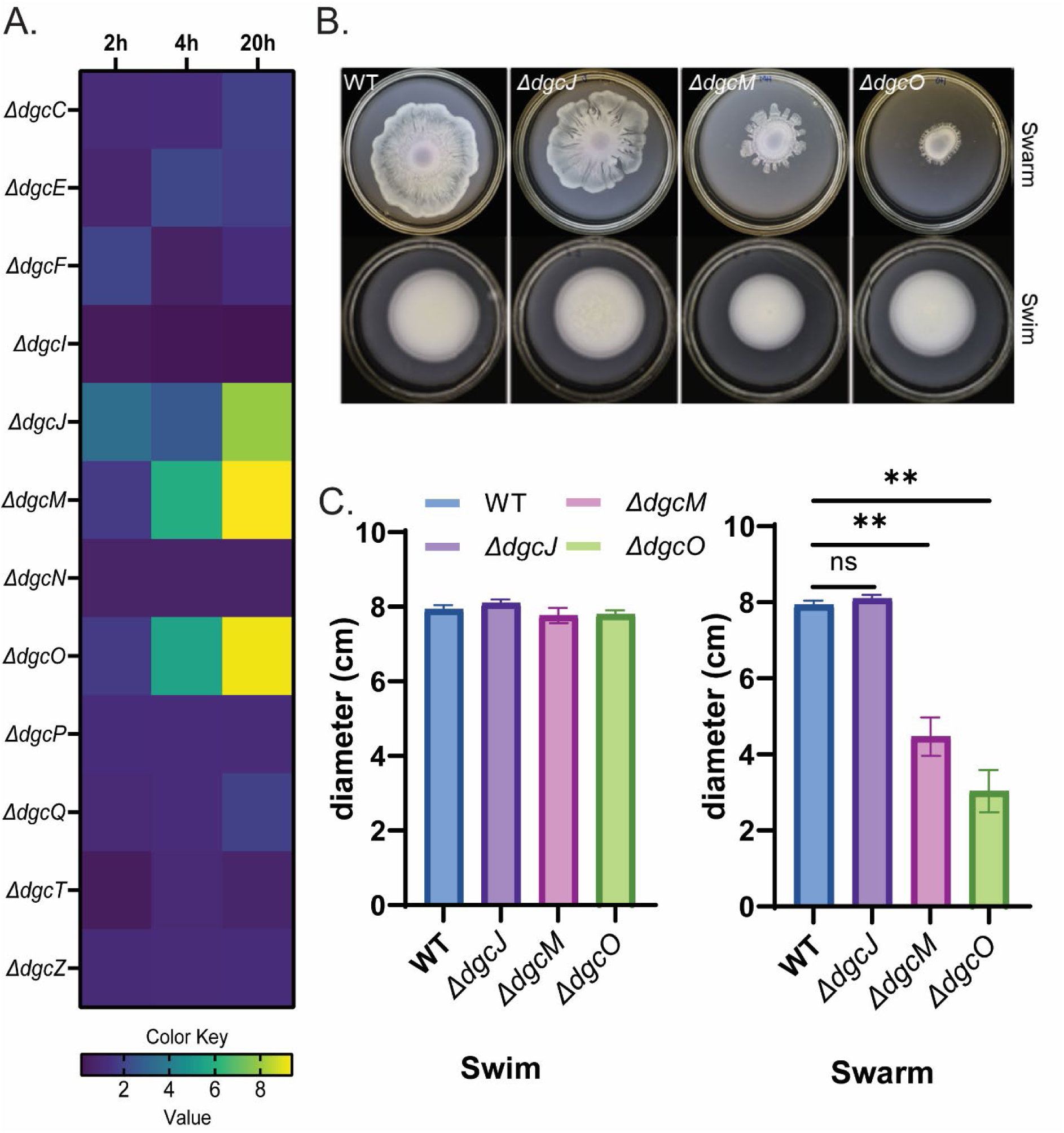
Identification of DGCs that contribute to swarming motility. (A) Comparison of fold changes in gene expression of WT *E. coli* DGCs during the time course of swarming. RNA-Seq data collected at 2, 4 and 20h were normalized to those from planktonic cultures (n=4). (B) Comparison of swimming (0.3% agar) and swarming (0.5% agar) motility of indicated DGC deletion mutants in the WT strain. Plates were incubated at 30℃ for 18h (n=3). (C) Plot of diameters across the zones of bacterial swimming or swarming shown in B. Calculated P values are indicated: *, <0.05, **, <0.01, or ***, <0.0001. NS, not statistically significant.

**Table 1.** c-di-GMP levels estimated from a riboswitch-based sensor.

| Strain | Relative c-di-GMP level to WT | Condition |
| --- | --- | --- |
| WT(MG1655)* | 1±0.008 | Swim |
| $\Delta pdeH$ | 2.88±0.11 | Swim |
| $\Delta dgcO$ | 0.82±0.03 | Swim |
| WT+pPdeH (10µM IPTG) | 0.75±0.04 | Swim |
| $\Delta dgcO$ + pDgcO (0% ara) | 1.28±0.37 | Swim |
| $\Delta dgcO$ + pDgcN (0% ara) | 1.5±0.09 | Swim |
| $\Delta dgcO$ + pDgcN+ (0.05% ara.) | 5±0.56 | Swim |
| WT(MG1655) | 1±0.093 | Swarm |
| $\Delta pdeH$ | 1.25±0.031 | Swarm |
| $\Delta dgcM$ | .89±0.0305 | Swarm |
| $\Delta dgcO$ | 0.57±0.048 | Swarm |
| WT+pPdeH (0µM IPTG) | 0.82±0.111 | Swarm |
| WT+pPdeH (10µM IPTG) | 0.67±0.18 | Swarm |
| WT+pPdeH (100µM IPTG) | 0.62±0.08 | Swarm |
| WT+pPdeH (1mM IPTG) | 0.21±0.04 | Swarm |
| $\Delta dgcO$ + pDgcO (0% ara) | 0.70±0.003 | Swarm |
| $\Delta dgcO$ + pDgcN (0% ara) | 1.48±0.013 | Swarm |
| $\Delta dgcO$ + pDgcO (0.05% ara) | 2.56±0.234 | Swarm |
| $\Delta dgcO$ + pDgcN+ (0.05% ara.) | 4.17±0.400 | Swarm |

\*Estimates of c-di-GMP levels from WT cells taken directly from a swim and swarm plates (see Methods) were set to 1 for comparison with all other strains and conditions tested. Based on reported data for planktonic *E. coli* [37], these values are roughly equivalent to 8pmol/mg protein. We are concerned here only with relative, not absolute, c-di-GMP values.

### c-di-GMP is required for swarming

Complementation of Δ*dgcO* with a plasmid encoding *dgcO* driven from an arabinose-inducible promoter (pDgcO) partially rescued the swarming defect without added inducer (i.e. leaky expression; Fig. 2A); addition of arabinose inhibited rescue, suggesting that a narrow window of c-di-GMP levels contributed by DgcO supported swarming. To test whether the Δ*dgcO* phenotype is related solely to alterations in c-di-GMP levels, we introduced into the mutant a plasmid encoding *dgcN* (_p_DgcN). Leaky expression of *dgcN* was sufficient to rescue swarming, while addition of inducer was inhibitory (Fig. 2A and Table 1). The swarming inhibition observed by induction of pDgcN in the Δ*dgcO* mutant (Fig. 2A) was relieved by deletion of *ycgR*, showing that the inhibition at the higher levels of c-di-GMP is due to interference with flagellar function as established in studies on swimming (Fig. S2). To confirm the assessment that a certain threshold level of c-di-GMP is required for swarming, we took an opposite approach, reducing c-di-GMP levels in the WT by introducing a plasmid expressing *pdeH* (pPdeH) under the control of an IPTG-inducible promoter. Even leaky expression had an inhibitory effect. Addition of as low a concentration as 10 μM IPTG reduced c-di-GMP levels below WT (Table 1), inhibiting swarming (Fig. 2B and 2C, left), but not swimming (Fig. 2C, right). c-di-GMP levels in all of these genetic backgrounds, in cells taken directly from the swarm and swim plates, are plotted in Fig. 2B,C and tabulated in Table 1. There was a correspondence between the measured c-di-GMP levels (Fig. 2D, left) and the extent of swarming (Fig. 2C, left), a correspondence that did not hold for swimming (compare 2C right with 2D right), suggesting that within the range tested, swarming is sensitive to c-di-GMP levels while swimming is not. We conclude that under our experimental conditions, c-di-GMP is necessary and supports optimal swarming within a narrow concentration range.

**Figure 2.**
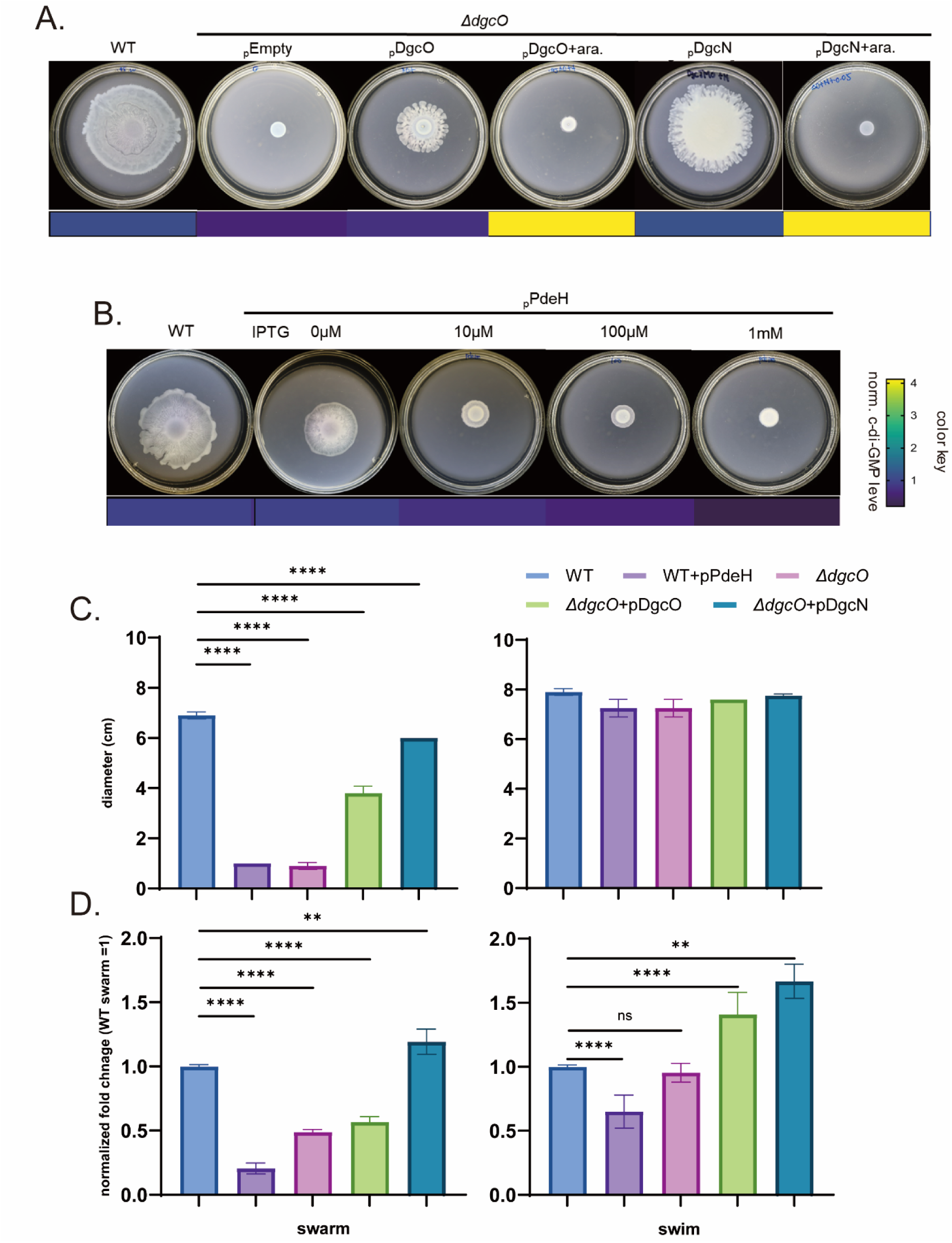
c-di-GMP is required for swarming. (A) Plasmids encoding *dgcO* (pDgcO), *dgcN* (pDgcN) or vector alone (pEmpty) under the control of pBAD promoter, were introduced to the *ΔdgcO* strain, and swarming compared to WT, with (+ara) or without addition of 0.05% arabinose (n=4). The color key beneath the plates corresponds to relative c-di-GMP levels measured using the riboswitch-based c-di-GMP sensor, shown in B as heatmap (see Table 1). (B) A plasmid encoding *pdeH* (pPdeH) driven from the T5-*lacO* promoter was introduced into WT and swarming monitored at indicated IPTG concentrations (n=2). C-di-GMP levels for all strains in this figure are found in Table1, represented here by a heat map beneath the plates. (C) Diameters of swim and swarm zones of strains indicated by a color key (n=4); pDgcO and pDgcN data are without added inducer, and pPdeH with 10μM IPTG. (D) Cells from swim/swarm plates in C were collected and c-di-GMP levels measured using the c-di-GMP biosensor. The values were normalized to the c-di-GMP levels in WT cells. Calculated *P* values are indicated: *, <0.05, **, <0.01, or ***, <0.0001. NS, not statistically significant. Color key as in C.

### Colanic acid is important for swarming

To investigate how c-di-GMP positively regulates swarming, we initially looked for suppressors that would restore swarming in a *ΔdgcO* strain, but no useful leads emerged. Given that the mutant strain could ‘move in place’ but not venture out (Movie S2), we wondered if the mutant might be defective in the production of polysaccharides that could contribute to surface wetting. We therefore examined the RNAseq data for changes in all exopolymers – LPS, CA, PGA, cellulose. Of these, several genes in the CA (especially *wcaF* and *wcaJ)* and cellulose biosynthetic pathways were seen to be upregulated (Fig. 3A). However, our WT MG1655 strain has a defect in cellulose production [40], so we focused on examining the contribution of CA. Using qRT-PCR, we confirmed that the expression levels of *wcaJ* were significantly lower in the *ΔdgcO* strain compared to the WT (Fig. 3B), suggesting a possible involvement of this DGC in CA synthesis.

**Figure 3.**
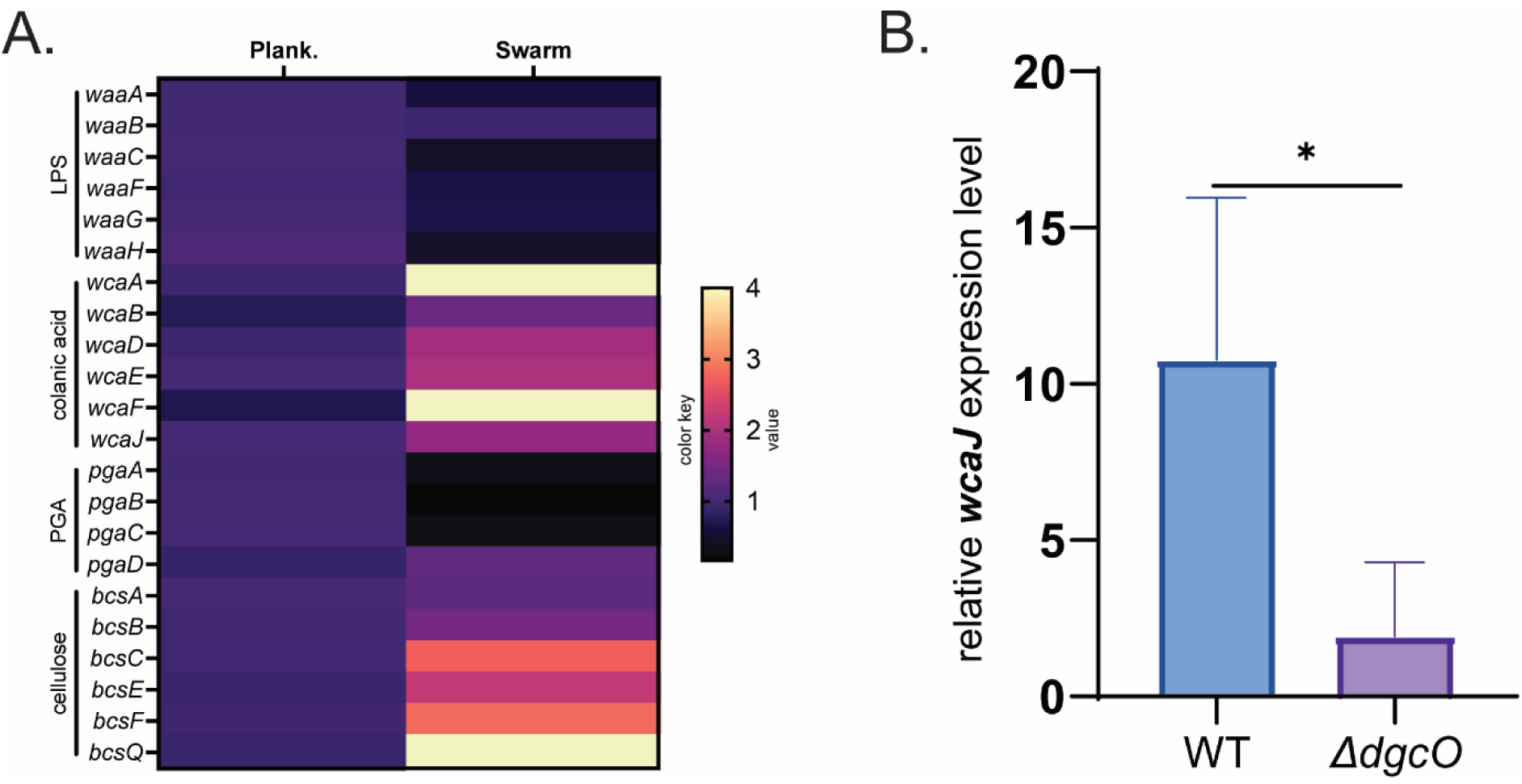
Exopolymer expression profiles during swarming. (A) Comparison of fold changes in gene expression of the four exopolymers of *E. coli* (LPS, CA, PGA, and cellulose). RNA-Seq data collected at 20h during the time course of swarming were normalized to those from planktonic cultures (n=4). (B) Comparison of *wcaJ* expression levels relative to the housekeeping gene *gyrA* in WT and *ΔdgcO,* using qRT-PCR from swarm cells collect at 20h (see Methods); (n=3).

To test the role of CA in swarming, we constructed two mutant strains: *ΔwcaJ* and *ΔwaaF*. *wcaJ* is part of a multi-gene operon that controls CA biosynthesis [41]; deletion of this gene interferes with CA synthesis but does not affect LPS synthesis. *waaF* belongs to a multi-gene operon controlling LPS biosynthesis [42]; deletion of this gene not only produces a defect in the LPS core, but also overproduces CA, possibly via the RcsCDB system [42]*. ΔwcaJ* was swarming defective (Fig. 4A). *ΔwaaF* could not be tested for swarming because flagella biosynthesis is inhibited in this strain [42]; however, we exploited its CA-overproduction phenotype to extract CA by established protocols (see Methods), adding it to the non-swarming *ΔwcaJ* directly on the swarm plate to test if external supplementation would rescue the defect (Fig. 4A). Through trial and error, we identified an amount of the extract that fully complemented the *ΔwcaJ* defect (Fig. 4A, +CA). It’s important to note that adding the same volume of water as a control was not effective, nor was a similar extract made from the *ΔwcaJ* strain (Fig. 4A).

**Figure 4.**
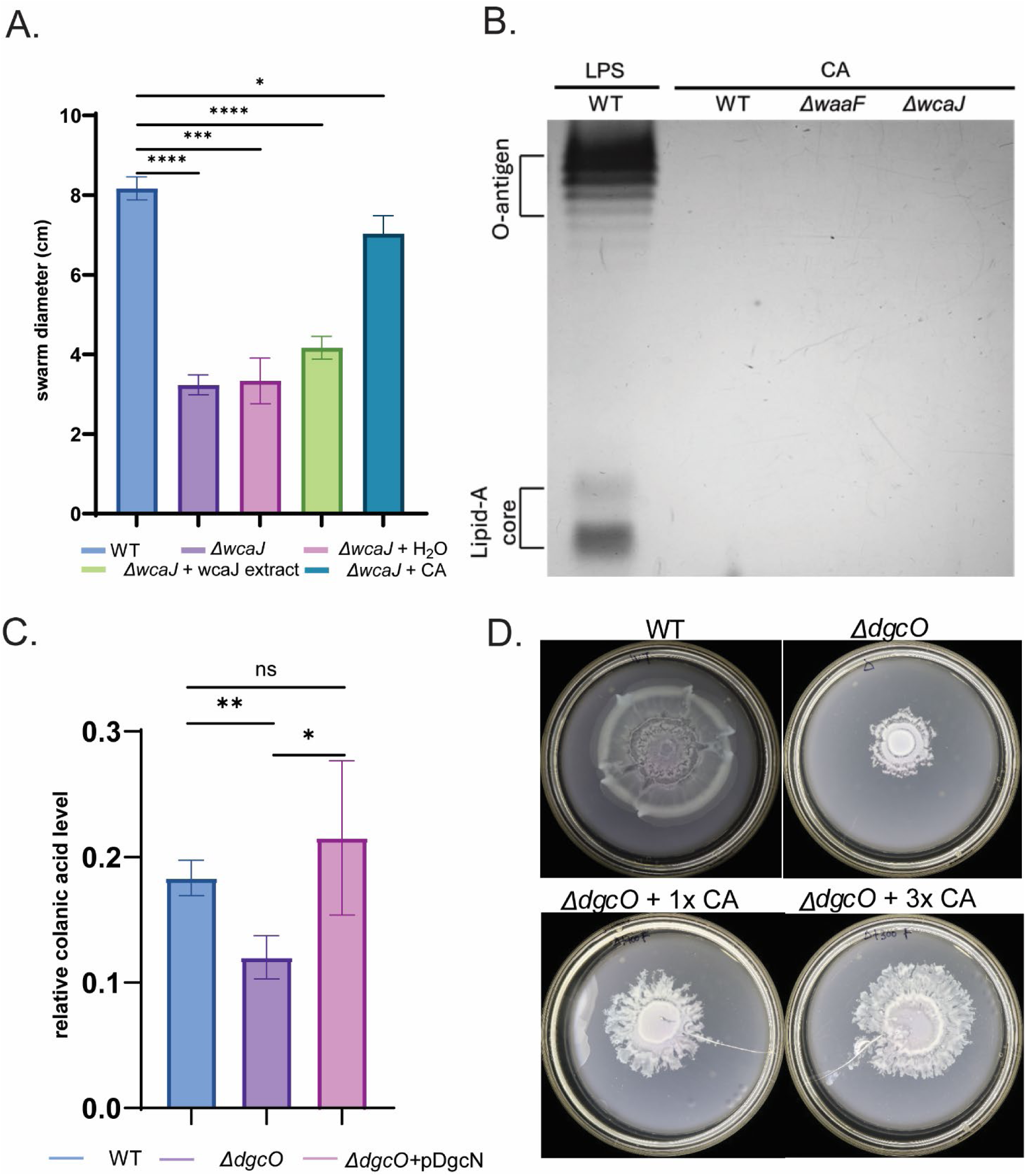
Colanic acid (CA) is required for swarming. (A) Swarming diameters at 20h were compared between WT, *ΔwcaJ* and *ΔwcaJ* supplemented with 100μl of either water, *ΔwcaJ* extract, or CA extract from *ΔwaaF* (n=3). (B) LPS extract from WT and CA extracts from *ΔwaaF* and *ΔwcaJ* strains were fractionated on SDS-PAGE gel, followed by silver staining (see Methods). (C) Comparison of relative CA levels between WT, *ΔdgcO* and *ΔdgcO* complemented with pDgcN (n=3). CA levels were measured as described under Methods. (D) Rescue of swarming in *ΔdgcO* by addition of indicated concentrations of CA extract from *ΔwaaF*.

To ensure that our CA extracts were not contaminated with LPS [41, 43], we compared them with LPS extracts from the WT strain [50] using a standard protocol [44]. Bands corresponding to LPS in WT extracts were absent in the CA extracts of *wcaJ* and *waaF* mutants (Fig. 4B). We conclude that CA and not LPS complements the *wcaJ* mutant of *E. coli* for swarming.

To determine if the swarming defect of *ΔdgcO* is due to lack of CA production, we first measured their levels in this strain (see Methods) (Fig. 4C). The *ΔdgcO* strain showed a significant decrease in CA production compared to the WT strain. Additionally, leaky expression from DgcN was sufficient to rescue CA levels in *ΔdgcO,* showing that c-di-GMP is required for maintaining CA production (Fig. 4C). To test if external addition of CA would rescue the *dgcO* defect, we added to it the CA extract from the *ΔwaaF* strain. We observed a positive correlation of swarming with the amount of CA added (Fig. 4D). Taken together, the data in Figure 4A-D allow us to conclude that DgcO is important for regulating the production of CA, which is a critical component for surface movement in *E. coli*. We have not investigated the nature of this regulation.

### Addition of glucose rescues *ΔdgcO* swarm defect

The availability of glucose has been reported to stimulate the Rcs regulon, which controls CA synthesis [41]. *E. coli* requires 0.5% glucose for optimal swarming, this dependency traditionally attributed to the need for maintaining the energy-intensive nature of swarming [45, 46]. However, examination of WT *E. coli* inoculated on swarm plates with either 0% glucose (no swarming) or 0.5% glucose, revealed that cells were similarly motile at the site of inoculation on both plates (Movie S3 and S4). Considering that addition of CA can rescue *ΔdgcO* cells from the swarm defect (Fig. 4D) and that glucose enhances CA production, we tested whether the addition of glucose could bypass the requirement for c-di-GMP signaling and enable the *ΔdgcO* strain to swarm. Swarming was monitored in four different strains - WT, *ΔdgcO, ΔwaaF*, and *ΔwcaJ* - on plates supplemented with either 0.5% or 1% glucose (w/v). *ΔdgcO* exhibited a full restoration of swarming at 1% glucose. In contrast, the *ΔwcaJ* strain showed only a slight increase in swarm diameter, likely due to the absence of a key component of the CA pathway in this mutant (Fig. 5A). (The lop-sided growth evident in the Δ*waaF* strain is due to the highly mucoid nature of excess CA.) These results were further verified by measuring relative CA levels across these plates (Fig. 5B), where addition of glucose increased the CA levels in both WT and *ΔdgcO,* but not in the *ΔwcaJ* strain. We conclude that CA is important for *E. coli* swarming.

**Figure 5.**
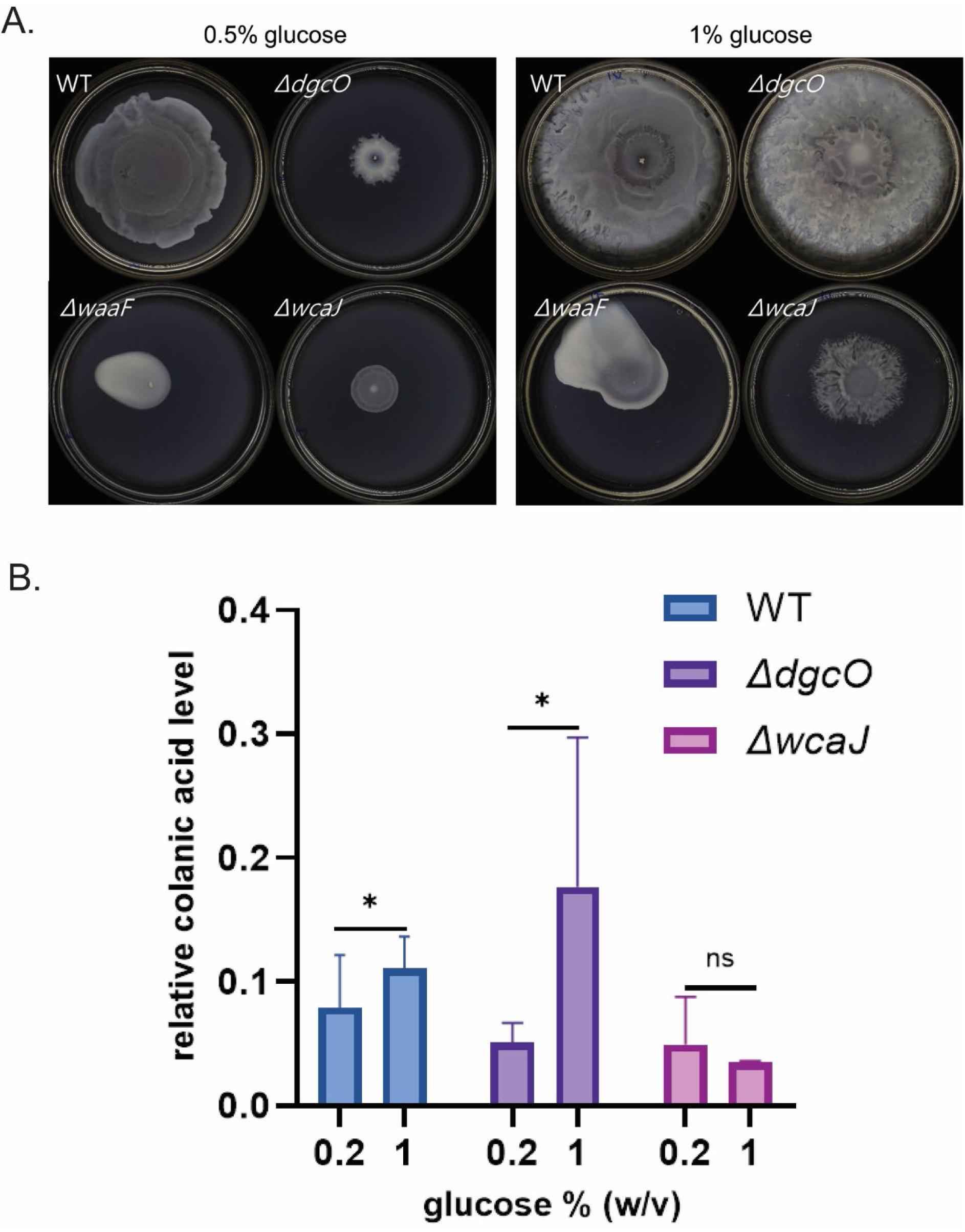
Addition of glucose rescues *ΔdgcO* swarm defect (A) Swarming recorded at 20h on plates supplied with two different glucose concentrations (0.5% or 1%) and inoculated with indicated strains. (B) Comparison of relative CA levels between WT, *ΔdgcO* and *ΔwcaJ* (n=3). CA levels were measured as described under Methods

### PGA is semi-essential for swarming, cellulose is not

In *E. coli*, both PGA and cellulose are regulated by c-di-GMP signaling [33, 37, 47–49], DgcO being implicated in PGA regulation [37]. To test the role of PGA in swarming, we constructed two mutant strains, *ΔpgaA* and *ΔpgaB*, that are crucial for PGA production. Neither strain was defective in swimming (Fig. S3A). Under standard conditions for preparing swarm plates, where we dry freshly poured plates for 1 h in a hood under laminar flow, these mutants were not significantly impaired for swarming compared to WT (Fig. S3B, top row). However, when the plates were dried for an extra 30 min (1.5h), a noticeable decrease in swarming was observed in the mutants compared to the WT. In *ΔpgaA*, this defect was overcome upon complementing *pgaA* from a plasmid (pPgaA), suggesting that PGA plays a niche role for *E. coli* motility on drier surfaces.

MG1655, the *E. coli* strain used in our study contains a mutation in the *bcsQ* gene in the cellulose synthesis pathway, that introduces an early stop codon, similar to many strains derived from the K-12 lineage [50]. This mutation results in reduced cellulose production compared to other *E. coli* strains, such as enteroaggregative (EAEC) that are known to form a robust biofilm [40]. To assess the role of cellulose in swarming motility, we reversed the premature stop codon by introducing a single mutation in codon 6 (TAG→TTG, A17T) of *bcsQ* to restore cellulose production as previously reported [40]. Indirect measurement of cellulose using crystal violet staining indicated a significant increase in staining (Fig. S3C). However, the reverted strain did no better than the original in swarming, even when plates were dried for an extra 30 minutes (Fig. S3D). From these experiments, we conclude that among the three polymers *E. coli* secretes, CA plays a major role, PGA a secondary role, and cellulose no role in swarming motility under our experimental conditions.

### Colanic acid acts as a surfactant

To understand how CA assists swarming motility, we had a choice between a wetting or a surfactant function. CA satisfies the former criterion, but we decided to evaluate the latter as well. To do so, we conducted a drop collapse test, a method commonly used to assay surfactants, where the curvature angle of the droplet is inversely correlated with its surfactant capacity [22]. CA extracts from WT, *ΔwcaJ*, and *ΔwaaF* strains propagated on swarm media were placed on a clean plastic surface, along with water as control. The measured curvature angle (see Methods) of the *ΔwcaJ* extract was comparable to that of water, whereas that of the *ΔwaaF* extract showed a significant decrease (Fig. 6A, B), demonstrating that CA indeed has surfactant properties. Next, we compared the curvature angle of these extracts immediately after being added to the surface of swarm media set with either Eiken or Fisher agar (outline marked with red-dotted lines) by capturing a close-up, top-view image of the drop (Fig. 6C, left). The larger the diameter of the drop base, the smaller the angle of contact [22] (Fig. 5C, right). Eiken agar showed a significantly lower contact angle compared to Fischer agar, indicating that the added water drop experiences lower surface tension on the former as surmised earlier ([24, 51]). The CA extract from Δ*waaF* placed on Eiken agar had the lowest contact angle of all. These results suggest that the addition of CA effectively lowers surface tension not only on plastic (Fig. 6A,B) but also on swarm agar plates. In summary, these data support our conclusion that CA has surfactant properties.

**Figure 6.**
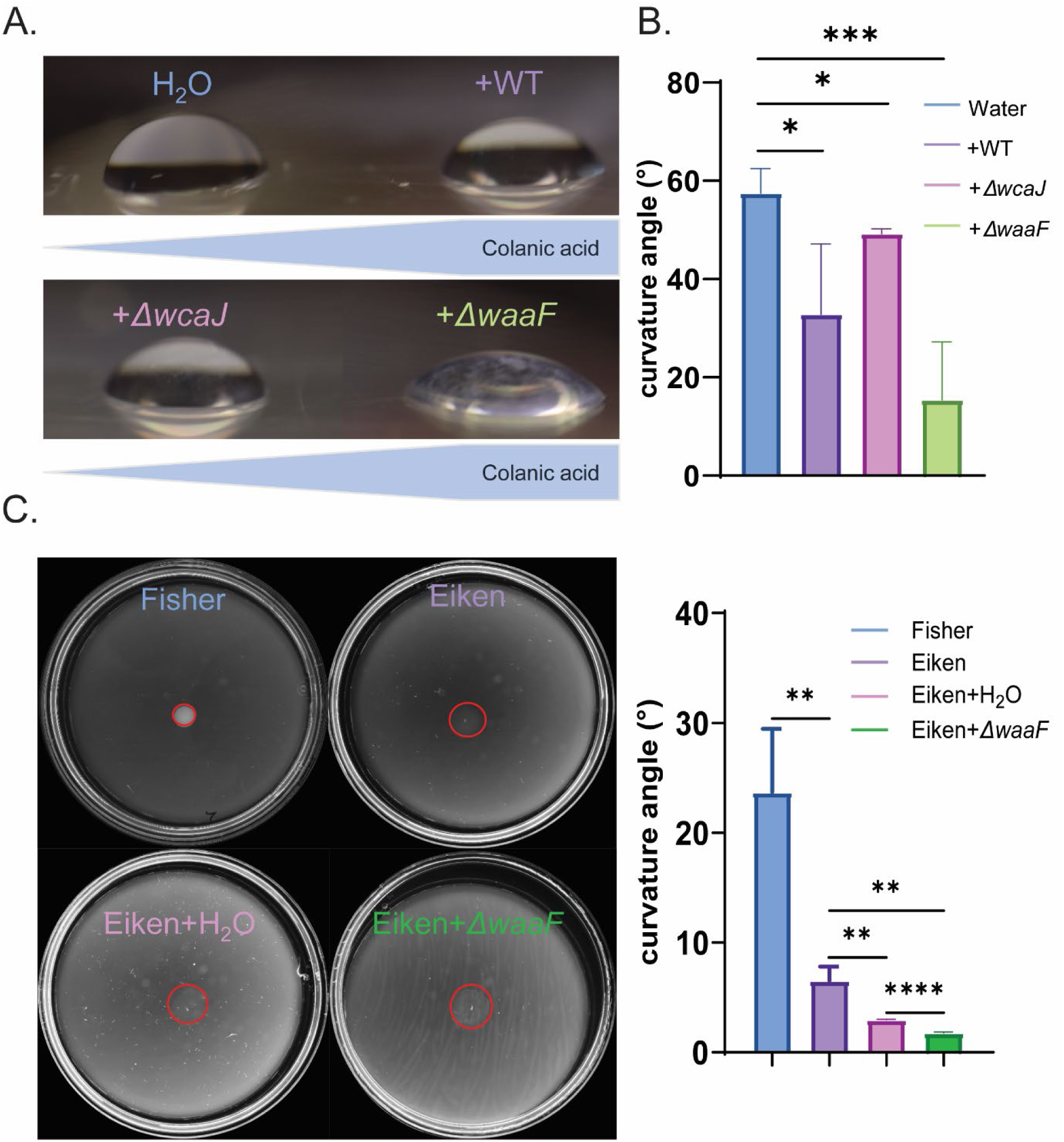
Colanic acid acts as a surfactant. (A) A side view of a 10-μl water drop or CA extracts in water from indicated strains, deposited on a plastic surface. Extracts with more CA are indicated in the blue ribbon below the images. The large contact angles could be measured directly from the image (see Methods). (B) A summary of measurements of contact angles from A. (C) Same as A-B, except water drops were placed directly on swarm media set with Fisher or Eiken agar (top row of plates), or water drops on Eiken agar compared with those of Δ*waaF* CA extract (bottom row of plates).

## Discussion

Studies on flagellar motility have thus far primarily focused on swimming populations to understand the role of c-di-GMP. This study explores how c-di-GMP influences swarming in *E. coli* and reports the following discoveries: 1. DgcJ/O/M are all upregulated during swarming (Fig. 1A). 2. DgcO was observed to play a major role, and DgcM a secondary role, in facilitating swarming, while no phenotype was evident for DgcJ under our experimental conditions (Fig. 1B,C). 3. c-di-GMP is required for swarming but enables this motility within a narrow range of c-di-GMP levels (Fig. 2 and Table 1). 4. DgcO promotes CA synthesis (Fig. 3). 5. External addition of CA to the swarming-defective *dgcO* mutant restores swarming (Fig. 4D). 6. CA has surfactant properties that are expected to aid swarming (Fig. 6A,B). 7. PGA, also known to be controlled by DgcO ([37]), contributes to swarming on drier surfaces (Fig. S3B). These findings are summarized in Figure 7.

**Figure 7.**
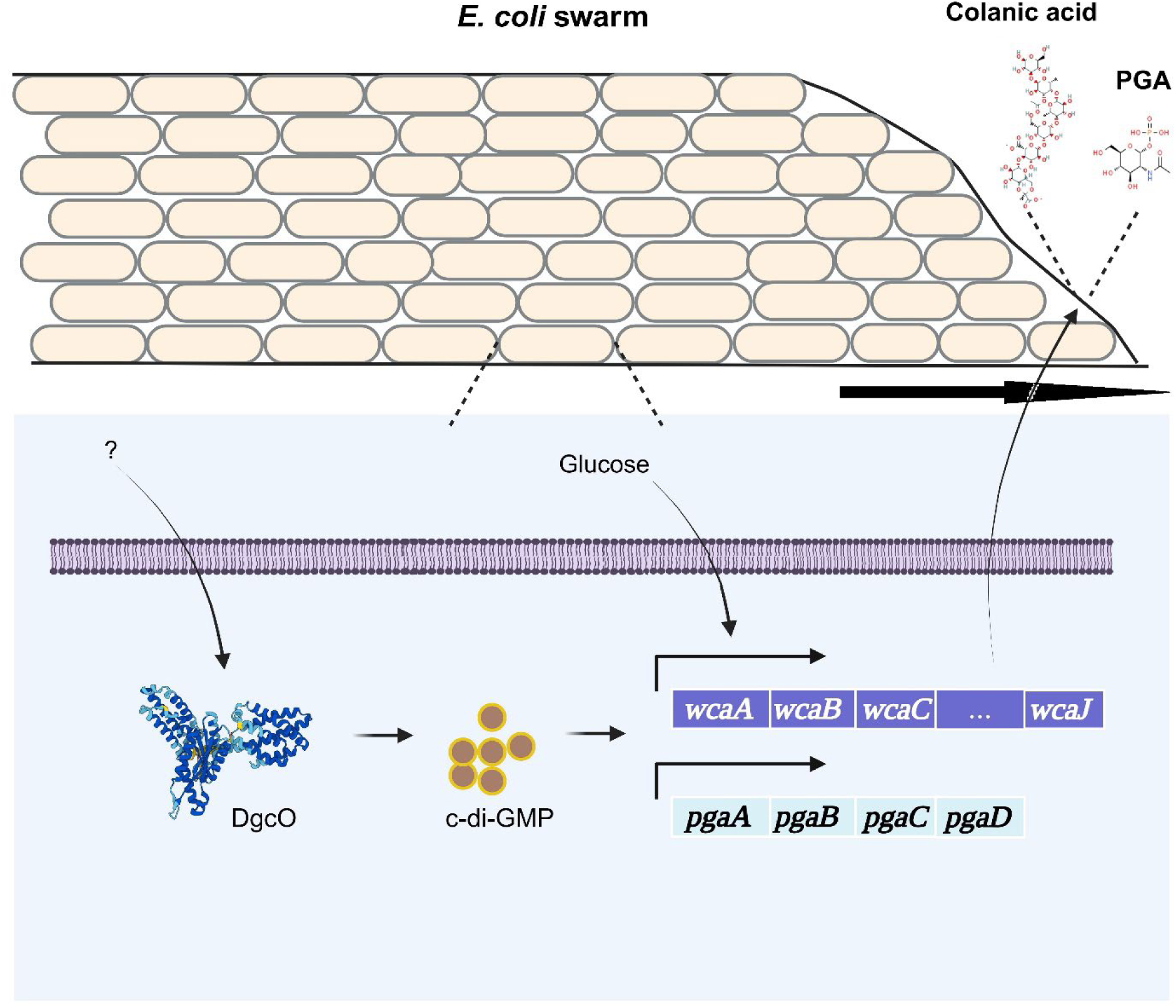
Summary of the role of DgcO in swarming. DgcO positively regulates swarming in *E. coli* by upregulating *wca* and *pga* operons engaged in the production of CA and PGA (this work and [37]. Glucose stimulates CA synthesis [41]. Depending on wet/dry surface conditions, these secreted exopolymers appear to function as wetting agents and surfactants to facilitate swarming.

A particularly gratifying aspect of this study is understanding why the *E. coli* strain we work with needs the special Eiken agar and addition of glucose, a requirement identified 30 years ago [24]. We show that plates set with Eiken agar have a lower surface tension compared to those with Fischer agar (not conducive to *E. coli* swarming) (Fig. 6C). We surmise that the role of added glucose is related to its known role in stimulating CA synthesis [41], thus additionally contributing to lowering surface tension.

Negatively charged polysaccharides have been hypothesized to act as wetting agents. That CA would also play such a role in *E. coli* is supported by a study that used osmolarity-sensing fluorescent liposomes to calculate that the wetting agent had to be a high molecular weight secreted substance [26]. While LPS would fit the bill, our study demonstrated that it is not a major player in our *E. coli* strain. We note that the robust swarmers *P. mirabilis* and *V. parahaemolyticus* both produce abundant capsular polysaccharides; for *P. mirabilis*, these are required for swarming [52, 53]. That such polysaccharides could also serve as surfactants, as shown here for CA, is a new realization.

While this study identified DgcO as specifically contributing to the regulation of CA synthesis, the ability of a heterologous DGC (DgcN) to complement the *ΔdgcO* defect (Fig. 2) does not exclude the possibility of a dedicated role for DgcO in the CA pathway, as observed for several other DGCs [54–57]. For instance in *P. aeruginosa*, the DGC GcbC directly interacts with its partner protein LapD, to achieve both specificity and maximal signaling output [57]. However, heterologous expression of DGC PA1107 can also induce biofilm formation through LapD [58]. Similarly, DGC LchD in *Lysobacter enzymogenes* forms a complex with its partner protein LchP to synthesize antibiotics; however, this can also be achieved by overexpression of another DGC [59]. In *E. coli*, a prior study showed an interaction between DgcO and DgcM, but that study only queried the interaction between their GGDEF domains [60]. Thus, more experiments are needed to address how DgcO influences CA synthesis.

When planktonic bacteria are transferred to a surface, they use a myriad surface and nutrient cues to change their physiology and adapt, as discussed extensively in a recent review [61]. c-di-GMP plays an important role in this process as demonstrated in several bacteria, particularly *P. aeruginosa* and *C. crescentus* [1, 3, 61]. A known nutrient cue for *E. coli* swarming is low intracellular iron [62]. In *P. aeruginosa* c-di-GMP levels are regulated by the amount of iron present [63]. The connection between iron and c-di-GMP remains to be elucidated in *E. coli*. We note that DgcO possesses a heme domain that interacts with oxygen [36, 64]. However, mutations in DgcO reported to interfere with heme binding did not reproduce the Δ*dgcO* swarming phenotype (data not shown).

Finally, while we have established conditions to observe *E. coli* swarming in the laboratory, the composition and texture of surfaces *E. coli* encounters in the environment must vary vastly. The role of DGCs or other enzymes contributing to facilitating collective motion may vary in importance accordingly.

## Acknowledgments

The c-di-GMP sensor was generously provided by Dr. Tai, Jung-Shen from Dr. Yan’s lab, Yale University. We thank Souvik Bhattacharyya for helpful comments during the course of this work, which was supported by NIH grant GM118085 to RMH.

## Methods

### Strains, growth conditions, genetic manipulations and motility assays

Strains and plasmids used in this study are listed in **Table S1**. The WT parent strain for *E. coli* was MG1655. Growth media and genetic manipulations have been described earlier [13]. Swarm plates were dried at room temperature (RT) for 1h inside Mystaire^®^ MY-PCR prep station laminar flow, prior to inoculation. CA or LPS extracts added were distributed across the surface by gentle shaking with beads (3mm) (CoilRollers Plating Beads, Novagen Co.).

### Determination of c-di-GMP levels by RFI

This method employs a riboswitch that specifically binds to c-di-GMP, causes a conformational change in the RNA structure that impacts downstream gene expression of RFP (Red Fluorescent Protein). A divergent constitutively active promoter controls CFP (Cyan Fluorescent Protein) expression and normalizes the data for cell number. Relative fluorescence intensity (RFI) of the two readouts at 574 nm (TurboRFP) and 489 nm (AmCyan) is a measure of relative c-GMP levels [65].

All c-di-GMP reporter strains were cultured overnight in gentamicin-containing medium at 37 °C with shaking at 200 rpm before being inoculated onto swim or swarm plates. The plates were incubated for 12 h (swim) or 20 h (swarm) and then stored at 4 °C for subsequent experiments. For swarm plate samples, cells were washed off the plates using PBS and resuspended to a final OD_600_ of 0.1. For swim plate samples, the central portion of the agar plate (diameter = 3 cm) was excised, transferred to a 50 mL tube, and washed with PBS. The tube was centrifuged at 1,000 × g for 10 minutes, the supernatant was discarded, and cells from the top of the agar were carefully collected using PBS and adjusted to a final OD_600_ of 0.1. Fluorescence spectra were measured using an RF-5301PC fluorescence spectrophotometer (Shimadzu, Kyoto, Japan). Samples were diluted with water to an OD_600_ of 0.1 before fluorescence measurement.

### Imaging phase contrast view of the edge of a swarm

A phase-contrast microscope (Olympus BX53) equipped with a 40× phase-contrast PH1 objective was used to observe the swarm front. A micro cover glass (18 × 18 mm, VWR) was carefully placed on top of the front and cell movement were captured using cellSens Imaging Software (Olympus Co.) at a rate of 10 frames per second, with a spatial resolution of 1004 × 997 pixels and a field of view measuring 120 × 120 μm² for up to 30s.

### RNA sequencing analysis of swarms

Analysis of *E. coli* swarm cells collected at 2, 4 and 20 h was performed by Marta Perez as part of another project. Swarm cells were harvested at 2, 4, and 20 h after inoculation. Cells were rinsed from the agar plates using a 2:1 mixture of RNAprotect Bacteria Reagent (Qiagen) and PBS and collected in 1.5 mL test tubes. A total volume of 1 mL was used to rinse all the cells present on each plate. Once collected, the bacterial suspension was vortexed for 5 seconds to mix thoroughly and then incubated at room temperature for 5 minutes. Cells were pelleted by centrifugation at 5000 x g for 10 minutes, and the supernatant was discarded. Total RNA was isolated from the cell samples treated with RNAprotect Bacteria Reagent using the RNeasy Mini Kit (Qiagen), following the enzymatic lysis and Proteinase K digestion protocol. A maximum of 6 × 10⁸ cells were processed per column. To prevent DNA contamination, on-column DNA digestion was performed using the RNase-free DNase Set (Qiagen), following the manufacturer’s instructions. RNA was eluted in 40 µL of RNase-free water and immediately kept cold.

RNA sequencing was outsourced to Novogene. RNA quality was assessed via visualization on agarose gel and determination of RNA concentration and RNA Integrity Number (RIN) using a Bioanalyzer (Agilent). The RNA quantity used for sequencing was at least 0.5 µg, with a minimum concentration of 10 ng/µL. The RIN values were ≥6 with a smooth baseline. OD_260/280_ and OD_260/230_ ratios were 2.0 or higher. Sequencing was performed on the Illumina NovaSeq 6000 system with a sequencing depth of 6.7 million reads per sample and a read length of 150 bp (paired-end).

### Quantitative reverse transcriptase PCR

*E. coli* cells grown on swarm plates (at 20h) were collected with PBS and resuspended in a final concentration of OD_600_=1. RNA was then purified using Qiagen RNeasy Mini kit (Qiagen co.) according to the manufacturer protocol. RNA concentration was determined using NanoDrop™ One (Thermo Scientific). RNA (1 μg) from each sample was subjected to 1-step qPCR using SuperScript III One-Step (Thermo Fisher Sci.), according to the manufacturer protocol. qRT-PCR reactions were performed in triplicates and fluorescence detection was performed using QuantStudio™ 7 Real-Time PCR (Thermo Fisher Sci.). RNA expression was normalized to the level of *gyrA*, a housekeeping gene control. The relative gene expression levels of *wcaJ* were calculated from cycle threshold (CT) values using the 2^−ΔC^ method, where ΔC = CT(*wcaJ*) – CT(*gyrA*) [66].

### Colanic acid extraction

Colanic acid (CA) was extracted and quantified by modification of the following method [67]: 50 µl of prepared extracts were then mixed with 4.5ml of H_2_SO_4_/H_2_O (6:1 v/v) and incubated at 100℃ for 20 min. The mixture was cooled to the RT and absorbances measured at 396 nm and 427 nm. 100µl of 1M cysteine hydrochloride (cys) were then added and the absorbances measured again at 396 nm and 427 nm. Final CA concentration was measured using the following equation.

[OD_396_ cys – OD_396_ pre-cys] – [OD_427_ cys – OD_427_ pre-cys]

### Extraction and silver staining of LPS

LPS was extracted by hot phenol-water method as described previously with some modifications [43]. To remove protein and nucleic acids from cell extracts, proteinase K (50 µg/mL) (Roche, Mannheim, Germany), RNase (40 µg/mL) (Roche, Mannheim, Germany), and DNase (20 µg/mL) (Roche, Mannheim, Germany) were added to the cell extract in the presence of 1 µL/mL 20% MgSO_4_ and incubated at 37°C overnight. An equal volume of hot (65°C) 90% phenol was added to the mixtures followed by vigorous shaking at 65°C for 15 min. Suspensions were then cooled, transferred to 1.5 mL polypropylene tubes and centrifuged at 8500×g for 15 min. Phenol phases were re-extracted by 300 µL distilled water. Sodium acetate at 0.5 M final concentration and 10 volumes of 95% ethanol were added to the extracts and samples were stored at −20°C overnight. After centrifugation, pellets were washed twice with 95% ethanol and resuspended in 120µl Laemmli Sample Buffer (33 mM Tris-HCl, pH 6.8, 1% SDS, 13.3% (w/v) glycerol, 0.005% bromophenol blue). Samples were then heated at 100℃ for 20 min. 10 µL of each sample (1×10^9^ cells) was separated on 15% SDS polyacrylamide gel with a 5% stacking gel at 100 mA for 1.5h. Silver and Coomassie blue staining of the gels was performed according to the standard protocols [68].

### Biofilm assay using Crystal Violet

Biofilm was quantified by using the following method [69].

### Contact angle measurement

Water or bacterial culture supernatant drops 10 μl in volume were placed on the outer side of the bottom part of 100 × 15-mm plastic Petri dishes (0875712; Fisher Scientific). The Petri dishes were washed with ethanol. For small angles of contact, where sideview images are not accurate, i.e. <30°, the diameter of each drop was determined by taking a close-up, top-view picture of the drop using a 24.2-megapixel camera (EOS R6 Mark II, Canon) with a 60-mm lens. Pictures were taken 5 min after drop deposition. The angle of contact, *θ* (°), between drop and surface was determined using a spherical-cap-shaped approximation for small angles (*θ* < 30°) through θ=720·Vπ2·a3 (where *V* is a set volume of the drop (10 *μ*l), and *a* is the radius (mm) of the circle formed by the drop base) as previously described [22].

## Supplementary Figures and Tables

**Figure S1.**
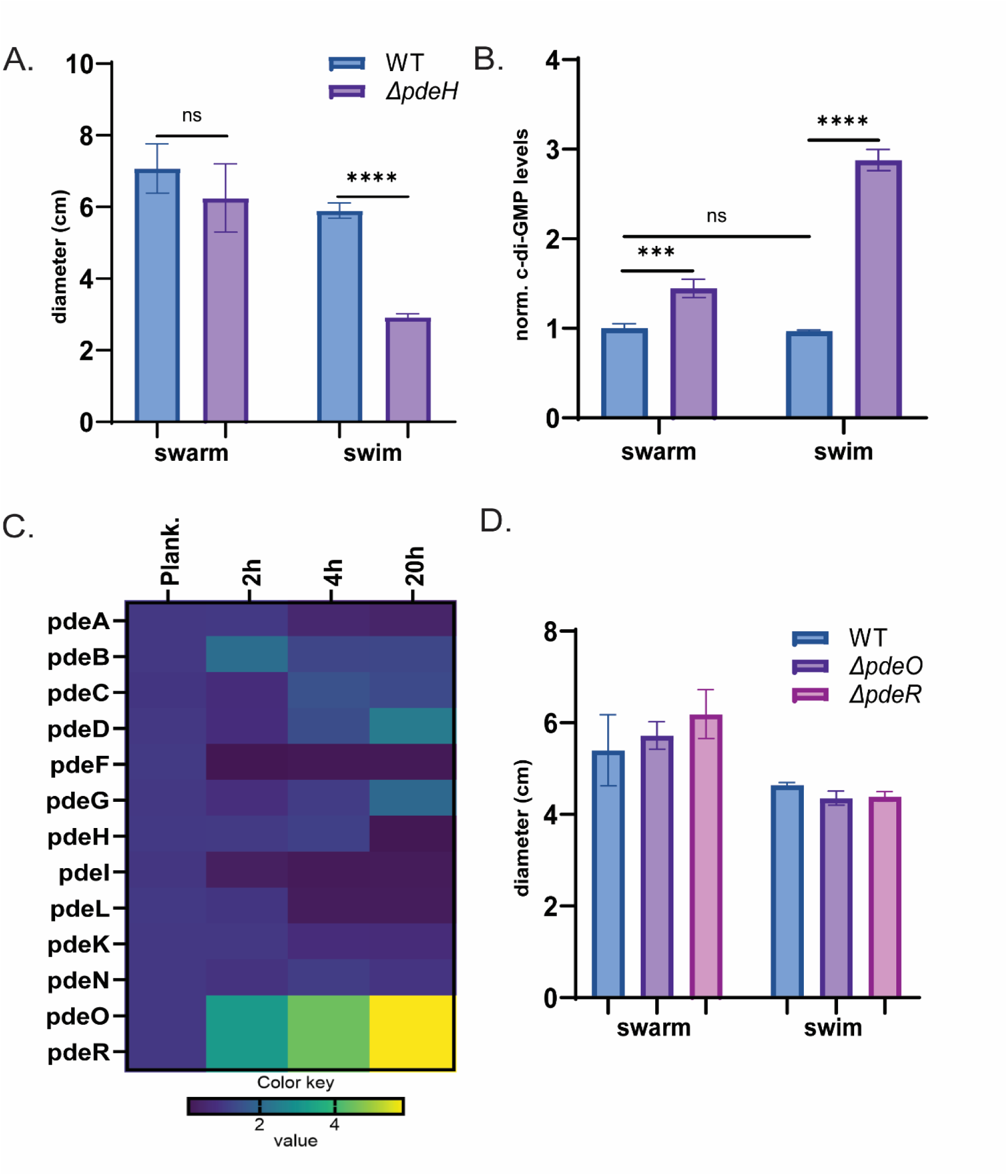
Differential c-di-GMP levels in swim and swarm cells. (A) Comparison of motility in swim (0.3% agar) or swarm (0.5% agar) plates of WT (MG1655) and Δ*pdeH* strains. Plates were incubated at 30℃ for 18h (n=3). (B) Measurement of c-di-GMP levels using a riboswitch-based c-di-GMP sensor (see Methods) in cells collected from swim and swarm plates shown in A, normalized to those of WT swim or swarm cells (see Table 1). Color coding of strains as in A. (C) Comparison of fold-changes in gene expression of all the *E. coli* PDEs during the time course of swarming. RNA-Seq data collected at 2, 4 and 20 h of swarming were normalized to those from planktonic cultures (n=4). (D) Swim and swarm motility comparison of WT, Δ*pdeO* and Δ*pdeR* strains as in A.

**Figure S2.**
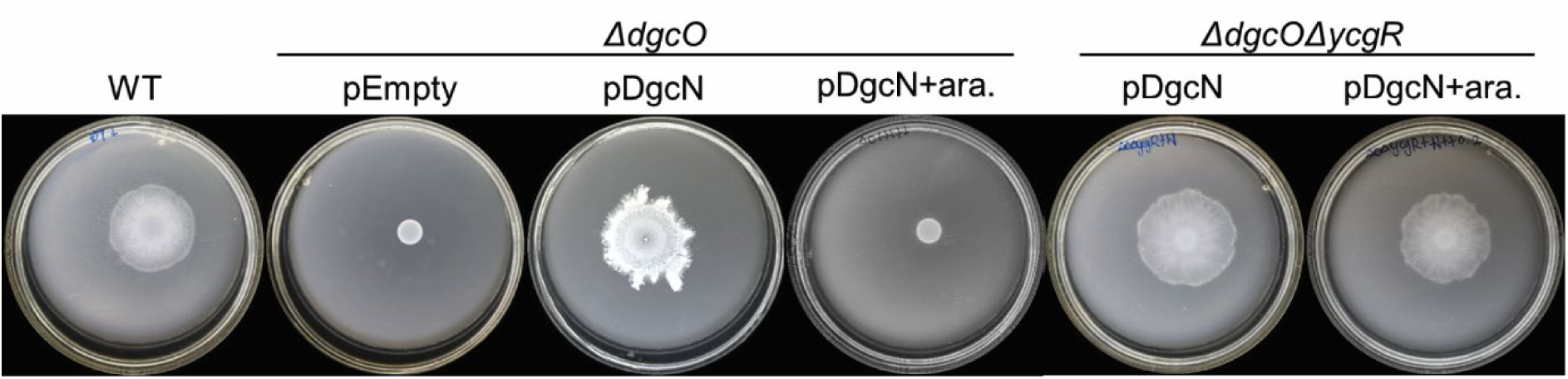
Swarming inhibition of *ΔdgcO* by pDgcN induction is through YcgR. Experimental setup as in Figure 2A except also monitored in a *ΔdgcOΔycgR* strain.

**Figure S3.**
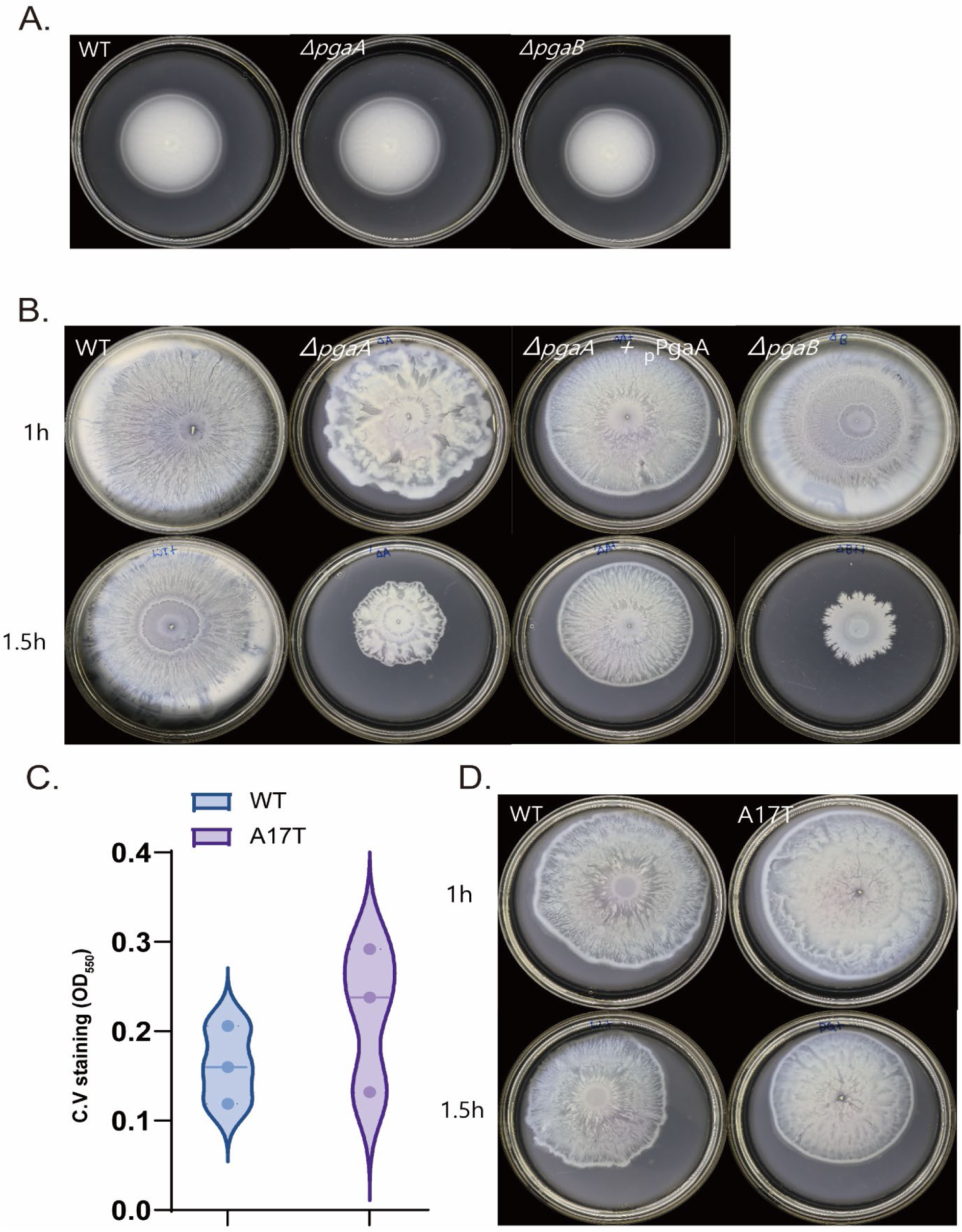
PGA, but not cellulose, assists swarming motility. (A-B) Comparison of swimming and swarming of indicated *pga* deletion mutants of MG1655. Plates were incubated at 30℃ for 18h (n=3) swarm or 37℃ for 12h (n=3) swim. Swarm plates were either dried under the hood for 1h (standard condition) or 1.5h. (C) Biofilm levels measured levels at OD_550nm_ by crystal violet staining of WT and *bcsQ*(A17T). (D) Swarming in indicated strains. Plates were dried for either 1 or 1.5h before cells were inoculated and incubated at 30℃ for 18h.

---

**Movie S1. Phase contrast view of the edge of a WT swarm**. Cells inoculated on swarm agar (see Methods) were viewed under phase contrast after 4h of incubation at 30°C, a time when movement is generally first observed. The movie is in real time, at 40x magnification.

**Movie S2. Phase contrast view of the edge of a Δ*dgcO* swarm**. Experimental protocol was as described in Movie S1.

**Movies S3 and S4. Movement at the edge WT *E. coli* swarm under different glucose concentrations**. Experimental conditions as described for Movie S1, except plates were supplemented with either 0% glucose (Movie S3) or 0.5% glucose (Movie S4).

**Table S1.**

| **Strains** | **Genotype or Description** | **Reference** |
| --- | --- | --- |
| MG1655 | K12 wild-type (WT) strain: F<sup>-</sup>, λ<sup>-</sup>, rph-1 | Laboratory Collection |
| JH702 | MG1655 $\Delta dgcM$ | This work |
| JH703 | MG1655 $\Delta dgcO$ | This work |
| JH1011 | MG1655 $\Delta dgcO$ + pBAD30_empty | This work |
| JH1012 | MG1655 $\Delta dgcO$ + pBAD30_DgcO | This work |
| JH1013 | MG1655 $\Delta dgcO$ + pBAD30_YflN | This work |
| JH706 | MG1655 $\Delta waaF$ | This work |
| JH707 | MG1655 $\Delta wcaJ$ | This work |
| JH711 | MG1655 + ASKA_pdeH | This work |
| JH1422 | MG1655 $\Delta dgcM$ + pBAD30_empty | This work |
| JH1423 | MG1655 $\Delta dgcM$ + pBAD30_DgcM | This work |
| JH1408 | MG1655 + cdiGsens | This work |
| JH1409 | MG1655 $\Delta dgcM$+ cdiGsens | This work |
| JH1410 | MG1655 $\Delta dgcO$+ cdiGsens | This work |
| JH1418 | MG1655 $\Delta pdeH$+ cdiGsens | This work |
| JH1105-1114 | MG1655 $\Delta dgcO$ suppressors | This work |
| JH1419 | MG1655 ASKA_PdeH+cdiG sens | This work |
| JH1201 | MG1655 $\Delta dgcO\Delta fhuA$ | This work |
| JH1202 | MG1655 $\Delta dgcO\Delta stfP$ | This work |
| JH1203 | MG1655 $\Delta dgcO\Delta yhbX$ | This work |
| JH1204 | MG1655 $\Delta dgcO\Delta fhuA$+cdiG sens | This work |
| **Plasmids** | **Expressed Proteins** |  |
| pKD4 | Kanamycin resistance gene template | [62] |
| pKD46 | λ Red Recombinase | [62] |
| pCP20 | FLP recombinase | [62] |
| pBAD30 | Cloning vector; pBAD; Amp<sup>R</sup> | [67] |
| pBAD30_YfiN | pBAD::YfiN | [67] |
| pBAD30_DgcO | pBAD::DgcO | This study |
| pBAD30_DgcM | pBAD::DgcM | This study |
| ASKA_PdeH | T5-*lacO* (pca24N)::PdeH | [68] |
| cdiG sens | pRP0122-*Pbe-amcyan_Bc4_turborfp* | [37] |

## References

1. Jenal, U., A. Reinders, and C. Lori, Cyclic di-GMP: second messenger extraordinaire. Nature Reviews Microbiology, 2017. 15(5): p. 271–284.

2. Povolotsky, T.L. and R. Hengge, ’Life-style’ control networks in Escherichia coli: signaling by the second messenger c-di-GMP. J Biotechnol, 2012. 160(1-2): p. 10–6.

3. Hengge, R., Principles of c-di-GMP signalling in bacteria. Nature Reviews Microbiology, 2009. 7(4): p. 263–273.

4. Hengge, R., et al., Systematic Nomenclature for GGDEF and EAL Domain-Containing Cyclic Di-GMP Turnover Proteins of Escherichia coli. Journal of bacteriology, 2015. 198(1): p. 7–11.

5. Boehm, A., et al., Second messenger-mediated adjustment of bacterial swimming velocity. Cell, 2010. 141(1): p. 107–16.

6. Pesavento, C., et al., Inverse regulatory coordination of motility and curli-mediated adhesion in Escherichia coli. Genes & development, 2008. 22(17): p. 2434–2446.

7. Sanchez-Torres, V., H. Hu, and T.K. Wood, GGDEF proteins YeaI, YedQ, and YfiN reduce early biofilm formation and swimming motility in Escherichia coli. Appl Microbiol Biotechnol, 2011. 90(2): p. 651–8.

8. Paul, K., et al., The c-di-GMP binding protein YcgR controls flagellar motor direction and speed to affect chemotaxis by a “backstop brake” mechanism. Mol Cell, 2010. 38(1): p. 128–39.

9. Fang, X. and M. Gomelsky, A post-translational, c-di-GMP-dependent mechanism regulating flagellar motility. Mol Microbiol, 2010. 76(5): p. 1295–305.

10. Girgis, H.S., et al., A comprehensive genetic characterization of bacterial motility. PLoS Genet, 2007. 3(9): p. 1644–60.

11. Han, Q., et al., Flagellar brake protein YcgR interacts with motor proteins MotA and FliG to regulate the flagellar rotation speed and direction. Front Microbiol, 2023. 14: p. 1159974.

12. Hou, Y.J., et al., Structural insights into the mechanism of c-di-GMP-bound YcgR regulating flagellar motility in Escherichia coli. J Biol Chem, 2020. 295(3): p. 808–821.

13. Nieto, V., et al., Under Elevated c-di-GMP in Escherichia coli, YcgR Alters Flagellar Motor Bias and Speed Sequentially, with Additional Negative Control of the Flagellar Regulon via the Adaptor Protein RssB. J Bacteriol, 2019. 202(1).

14. Valentini, M. and A. Filloux, Multiple Roles of c-di-GMP Signaling in Bacterial Pathogenesis. Annu Rev Microbiol, 2019. 73: p. 387–406.

15. Limoli, D.H., C.J. Jones, and D.J. Wozniak, Bacterial Extracellular Polysaccharides in Biofilm Formation and Function. Microbiol Spectr, 2015. 3(3).

16. Tuson, H.H. and D.B. Weibel, Bacteria-surface interactions. Soft Matter, 2013. 9(18): p. 4368–4380.

17. Partridge, J.D. and R.M. Harshey, Swarming: flexible roaming plans. J Bacteriol, 2013. 195(5): p. 909–18.

18. Kearns, D.B., A field guide to bacterial swarming motility. Nat Rev Microbiol, 2010. 8(9): p. 634–44.

19. Partridge, J.D. and R.M. Harshey, Flagellar protein FliL: A many-splendored thing. Mol Microbiol, 2024. 122(4): p. 447–454.

20. Be’er, A. and G. Ariel, A statistical physics view of swarming bacteria. Mov Ecol, 2019. 7: p. 9.

21. Chen, B.G., L. Turner, and H.C. Berg, The wetting agent required for swarming in Salmonella enterica serovar typhimurium is not a surfactant. J Bacteriol, 2007. 189(23): p. 8750–3.

22. Be’er, A. and Rasika M. Harshey, Collective Motion of Surfactant-Producing Bacteria Imparts Superdiffusivity to Their Upper Surface. Biophysical Journal, 2011. 101(5): p. 1017–1024.

23. Partridge, J.D. and R.M. Harshey, More than motility: Salmonella flagella contribute to overriding friction and facilitating colony hydration during swarming. J Bacteriol, 2013. 195(5): p. 919–29.

24. Harshey, R.M. and T. Matsuyama, Dimorphic transition in Escherichia coli and Salmonella typhimurium: surface-induced differentiation into hyperflagellate swarmer cells. Proc Natl Acad Sci U S A, 1994. 91(18): p. 8631–5.

25. Harshey, R.M., Swarming Adventures., in The lure of bacterial genetics: a tribute to John Roth, Kelly T. Hughes, Stanley Maloy, Josep Casadesús, Editors. 2010, American Society for Microbiology: Washington, D.C. p. 163–72.

26. Toguchi, A., et al., Genetics of swarming motility in Salmonella enterica serovar Typhimurium: critical role for lipopolysaccharide. Journal of bacteriology, 2000. 182(22): p. 6308–6321.

27. Ping, L., et al., Osmotic pressure in a bacterial swarm. Biophys J, 2014. 107(4): p. 871–8.

28. Wang, Q., et al., Sensing wetness: a new role for the bacterial flagellum. Embo j, 2005. 24(11): p. 2034–42.

29. Mariconda, S., Q. Wang, and R.M. Harshey, A mechanical role for the chemotaxis system in swarming motility. Mol Microbiol, 2006. 60(6): p. 1590–602.

30. Wang, Q., et al., Gene expression patterns during swarming in Salmonella typhimurium: genes specific to surface growth and putative new motility and pathogenicity genes. Mol Microbiol, 2004. 52(1): p. 169–87.

31. Junkermeier, E.H. and R. Hengge, A Novel Locally c-di-GMP-Controlled Exopolysaccharide Synthase Required for Bacteriophage N4 Infection of Escherichia coli. mBio, 2021. 12(6): p. e0324921.

32. Sellner, B., et al., A New Sugar for an Old Phage: a c-di-GMP-Dependent Polysaccharide Pathway Sensitizes Escherichia coli for Bacteriophage Infection. mBio, 2021. 12(6): p. e0324621.

33. Junkermeier, E.H. and R. Hengge, Local signaling enhances output specificity of bacterial c-di-GMP signaling networks. Microlife, 2023. 4: p. uqad026.

34. Lindenberg, S., et al., The EAL domain protein YciR acts as a trigger enzyme in a c-di-GMP signalling cascade in E. coli biofilm control. Embo j, 2013. 32(14): p. 2001–14.

35. Serra, D.O. and R. Hengge, A c-di-GMP-Based Switch Controls Local Heterogeneity of Extracellular Matrix Synthesis which Is Crucial for Integrity and Morphogenesis of Escherichia coli Macrocolony Biofilms. Journal of Molecular Biology, 2019. 431(23): p. 4775–4793.

36. Tuckerman, J.R., et al., An oxygen-sensing diguanylate cyclase and phosphodiesterase couple for c-di-GMP control. Biochemistry, 2009. 48(41): p. 9764–74.

37. Tagliabue, L., et al., The diguanylate cyclase YddV controls production of the exopolysaccharide poly-N-acetylglucosamine (PNAG) through regulation of the PNAG biosynthetic pgaABCD operon. Microbiology (Reading), 2010. 156(Pt 10): p. 2901–2911.

38. Bhattacharyya, S., et al., Efflux-linked accelerated evolution of antibiotic resistance at a population edge. Molecular Cell, 2022. 82(22): p. 4368–4385.e6.

39. Zhou, H., et al., Characterization of a natural triple-tandem c-di-GMP riboswitch and application of the riboswitch-based dual-fluorescence reporter. Sci Rep, 2016. 6: p. 20871.

40. Serra, D.O., A.M. Richter, and R. Hengge, Cellulose as an architectural element in spatially structured Escherichia coli biofilms. J Bacteriol, 2013. 195(24): p. 5540–54.

41. Wang, C., et al., Colanic acid biosynthesis in Escherichia coli is dependent on lipopolysaccharide structure and glucose availability. Microbiol Res, 2020. 239: p. 126527.

42. Wang, Z., et al., Deletion of the genes waaC, waaF, or waaG in Escherichia coli W3110 disables the flagella biosynthesis. J Basic Microbiol, 2016. 56(9): p. 1021–35.

43. Rezania, S., et al., Extraction, Purification and Characterization of Lipopolysaccharide from Escherichia coli and Salmonella typhi. Avicenna J Med Biotechnol, 2011. 3(1): p. 3–9.

44. Tan, L. and P.S. Grewal, Comparison of two silver staining techniques for detecting lipopolysaccharides in polyacrylamide gels. J Clin Microbiol, 2002. 40(11): p. 4372–4.

45. Kim, W. and M.G. Surette, Metabolic differentiation in actively swarming Salmonella. Mol Microbiol, 2004. 54(3): p. 702–14.

46. Inoue, T., et al., Genome-wide screening of genes required for swarming motility in Escherichia coli K-12. J Bacteriol, 2007. 189(3): p. 950–7.

47. Wang, X., J.F. Preston, 3rd, and T. Romeo, *The pgaABCD locus of Escherichia coli promotes the synthesis of a polysaccharide adhesin required for biofilm formation*. J Bacteriol, 2004. 186(9): p. 2724–34.

48. Hammar, M., et al., Expression of two csg operons is required for production of fibronectin- and congo red-binding curli polymers in Escherichia coli K-12. Mol Microbiol, 1995. 18(4): p. 661–70.

49. Barnhart, M.M. and M.R. Chapman, Curli biogenesis and function. Annu Rev Microbiol, 2006. 60: p. 131–47.

50. Serra, D.O., G. Klauck, and R. Hengge, Vertical stratification of matrix production is essential for physical integrity and architecture of macrocolony biofilms of Escherichia coli. Environ Microbiol, 2015. 17(12): p. 5073–88.

51. Berg, H.C., Swarming Motility: It Better Be Wet. Current Biology, 2005. 15(15): p. R599–R600.

52. Rahman, M.M., et al., The structure of the colony migration factor from pathogenic Proteus mirabilis. A capsular polysaccharide that facilitates swarming. J Biol Chem, 1999. 274(33): p. 22993–8.

53. Boles, B.R. and L.L. McCarter, Vibrio parahaemolyticus scrABC, a novel operon affecting swarming and capsular polysaccharide regulation. J Bacteriol, 2002. 184(21): p. 5946–54.

54. Chen, G., et al., Structural basis for diguanylate cyclase activation by its binding partner in Pseudomonas aeruginosa. Elife, 2021. 10.

55. Xu, Z., et al., Interplay between the bacterial protein deacetylase CobB and the second messenger c-di-GMP. Embo j, 2019. 38(18): p. e100948.

56. Chen, G., et al., The SiaA/B/C/D signaling network regulates biofilm formation in Pseudomonas aeruginosa. The EMBO Journal, 2020. 39(6): p. e103412.

57. Dahlstrom Kurt, M., et al., Contribution of Physical Interactions to Signaling Specificity between a Diguanylate Cyclase and Its Effector. mBio, 2015. 6(6): p. 10.1128/mbio.01978-15.

58. Newell, P.D., R.D. Monds, and G.A. O’Toole, LapD is a bis-(3’,5’)-cyclic dimeric GMP-binding protein that regulates surface attachment by Pseudomonas fluorescens Pf0-1. Proc Natl Acad Sci U S A, 2009. 106(9): p. 3461–6.

59. Xu, G., et al., Diguanylate Cyclase and Phosphodiesterase Interact To Maintain the Specificity of Cyclic di-GMP Signaling in the Regulation of Antibiotic Synthesis in Lysobacter enzymogenes. Applied and Environmental Microbiology, 2022. 88(2): p. e01895–21.

60. Sarenko, O., et al., More than Enzymes That Make or Break Cyclic Di-GMP—Local Signaling in the Interactome of GGDEF/EAL Domain Proteins of Escherichia coli. mBio, 2017. 8(5): p. e01639–17.

61. Laventie, B.J. and U. Jenal, Surface Sensing and Adaptation in Bacteria. Annu Rev Microbiol, 2020. 74: p. 735–760.

62. Bhattacharyya, S., et al., A heritable iron memory enables decision-making in Escherichia coli. Proc Natl Acad Sci U S A, 2023. 120(48): p. e2309082120.

63. Zhan, X., et al., A c-di-GMP signaling module controls responses to iron in Pseudomonas aeruginosa. Nat Commun, 2024. 15(1): p. 1860.

64. Kitanishi, K., et al., Important roles of Tyr43 at the putative heme distal side in the oxygen recognition and stability of the Fe(II)-O2 complex of YddV, a globin-coupled heme-based oxygen sensor diguanylate cyclase. Biochemistry, 2010. 49(49): p. 10381–93.

65. Zhou, H., et al., Characterization of a natural triple-tandem c-di-GMP riboswitch and application of the riboswitch-based dual-fluorescence reporter. Scientific Reports, 2016. 6(1): p. 20871.

66. Livak, K.J. and T.D. Schmittgen, Analysis of relative gene expression data using real-time quantitative PCR and the 2(-Delta Delta C(T)) Method. Methods, 2001. 25(4): p. 402–8.

67. Obadia, B., et al., Influence of tyrosine-kinase Wzc activity on colanic acid production in Escherichia coli K12 cells. J Mol Biol, 2007. 367(1): p. 42–53.

68. Li, H. and M. Benghezal, Crude Preparation of Lipopolysaccharide from Helicobacter pylori for Silver Staining and Western Blot. Bio-protocol, 2017. 7(20): p. e2585.

69. O’Toole, G.A., Microtiter dish biofilm formation assay. J Vis Exp, 2011(47).

